# PyMouse Lifter: Real Time 3-D Pose Estimation for Mice with Only 2-D Annotation Via Data Synthesis

**DOI:** 10.1101/2025.08.02.668290

**Authors:** Haozong Zeng, Hao Hu, Tony Fong, Ami Citri, Timothy H. Murphy

**Affiliations:** University of British Columbia, Department of Psychiatry, Faculty of Medicine, Vancouver, British Columbia, Canada; Djavad Mowafaghian Centre for Brain Health, University of British Columbia, Vancouver, British Columbia, Canada; The Edmond & Lily Safra Center for Brain Sciences, The Hebrew University of Jerusalem, Israel; The Alexander Silberman Institute for Life Science, The Hebrew University of Jerusalem, Israel; Program in Child and Brain Development, Canadian Institute for Advanced Research, MaRS Centre, Canada

**Keywords:** 3-D capture, pose estimation, synthetic data, automation

## Abstract

Neural-network-based pose estimation models have become increasingly popular for quantitative analysis of mouse behavior, yet most recordings still use a single 2-D camera view and therefore lack the depth cues needed for accurate 3-D kinematics. Existing open-source 3-D mouse datasets for training deep-learning models cover only a narrow range of environments and do not generalize well to various laboratory settings. To overcome these limitations, we introduce **PyMouse Lifter**, a pipeline that automatically reconstructs 3-D mouse poses from ordinary 2-D top-view videos with minimal manual 2-D annotation. PyMouse Lifter combines (i) an anatomically realistic 3-D mouse model for automated data synthesis, (ii) a monocular depth estimation model, and (iii) a 2-D key-point estimation model, enabling accurate 3-D reconstruction (model-based 3D inference) in virtually any open-field arena without using depth or multiple camera views for reconstruction. We validate the system on multiple datasets against depth-camera ground truth and show that the lifted 3D trajectories yield improved behavior classification over 2-D data and can be implemented in real time.

## 1. Introduction

Mice serve as a central model in neuroscience and preclinical investigation. However, compared to humans, their small size and highly flexible bodies pose unique challenges for motion capture. Wearable markers are impractical, and although multi-camera or depth-camera systems are effective, they require specialized hardware setups and extensive labeling efforts. Consequently, 2-D pose estimation solutions such as DeepLabCut (Mathis et al., 2018), SLEAP (Pereira et al., 2022), and YOLO-based (Jocher & Qiu, 2024) methods (Hatton-Jones et al., 2021; Sun et al., 2021) have become standard for tracking rodent movements in two dimensions. However, these approaches often fail to capture the richer information afforded by full 3-D kinematics (Dunn et al., 2021; Karashchuk et al., 2021).

Recent methods for single-view 3-D pose estimation in animals (Gharagozloo et al., 2021; Hu et al., 2023) typically rely on supervised training with large amounts of labeled 3-D data, thereby limiting their applicability in diverse environments where such data are unavailable or prohibitively difficult to obtain. Yet, many downstream behavior detection methods—such as the Mouse Action Recognition System (MARS) (Segalin et al., 2021), MoSeq (Wiltschko et al., 2020), and keypoint-MoSeq(Weinreb et al., 2024)—rely on these object detection, depth and/or keypoint-based pose estimation results. Recently, pre-trained monocular depth estimation (MDE) models—such as MiDaS (Ranftl et al., 2022), ZoeDepth (Bhat et al., 2023), and Depth Anything (Yang et al., 2024)—have made significant strides in handling previously unseen scenarios. However, their direct application to rodent experiments typically requires specialized fine-tuning with metric depth data, which might be difficult to obtain. Research has shown that a promising strategy to deal with the absence of training data—common in human pose estimation—is to leverage synthetic datasets generated from high-fidelity 3-D models (Loper et al., 2015; Zuffi et al., 2017). These datasets provide an unlimited supply of labeled images free from annotator bias and can span numerous environmental and lighting conditions (Li et al., 2020; Yan et al., 2021). However, for mice, the lack of robust 3-D art assets and the limited commercial interest outside biomedical research impede the development of large-scale synthetic datasets. Bolaños et al. (2021) demonstrated the viability of using a 3-D mouse model and generative adversarial networks for style transfer, but generating the key-framed poses and aligning real and synthetic data remain tedious tasks requiring 3-D animation expertise.

In this study we present PyMouse Lifter, a synthetic-data pipeline that automatically turns ordinary single-camera recordings into a rich source of 3-D information, thereby enabling accurate pose estimation and behavior analysis when only 2-D single view footage is available. From a small set of manually annotated frames and a pre-trained monocular-depth network, the pipeline (Fig. 1A) auto-generates pseudo-labels—2-D keypoints and coarse depth maps. These pseudo labels drive a biomechanically constrained 3-D mouse model, which refines the poses and produces photorealistic renders with anatomically-localized keypoints and depth maps. A style-transfer stage then aligns the renders to laboratory lighting and texture. We demonstrate that models trained on this synthetic data can support real-time behaviour classification (Fig. 1B), extending advanced 3-D behavioral annotation to any lab equipped with a single webcam.

**Figure 1.**
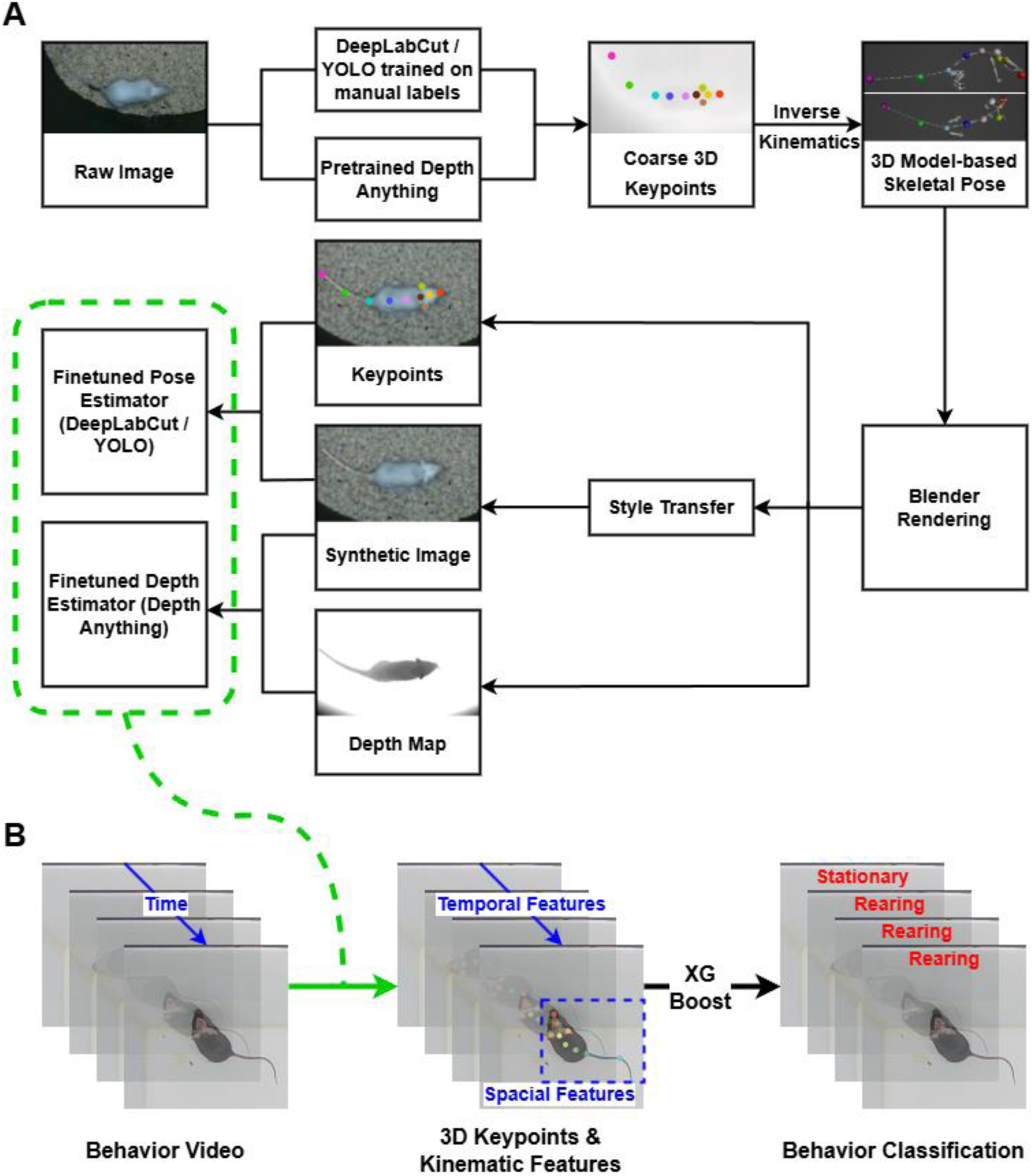
Overall workflow of PyMouse Lifter for achieving 3-D estimation from only 2-D images and minimal annotations by leveraging synthetic labelled data. (A) Simplified pipeline of creating synthetic training data using 2-D images captured without any 3-D capture hardware, then fine-tuning deep learning models to achieve 3-D estimation. (B) Schematic of the inference stage: the fine-tuned deep-learning models accept raw top-view RGB frames as input and directly output frame-wise 3-D keypoint coordinates, which are subsequently used for downstream behavioral-kinematic analysis.

## 2. Methods

### Animals

Behavioral recordings were obtained from the following cohorts: (i) 16 male C57BL/6 mice, 3–7 months old, with or without cranial windows as previously described (Fong et al., 2023); (ii) 2 female C57BL/6 mice, 3 months old; (iii) 2 FVB/N mice, 5 months old; and (iv) 2 male C57BL/6 mice, 13 months old. All animals were group-housed in standard plastic cages under a 12 h/12 h light–dark cycle (lights on at 07:00). Experiments were approved by the University of British Columbia Animal Care Committee and carried out in accordance with national guidelines.

### Experiments and Data Acquisition

Open-field videos with 3-D ground truth were recorded using an Orbbec Gemini 335L structured-light depth camera mounted above a 32 cm × 32 cm open-field arena (Fig. 2), following the protocol described in PyMouseTracks (Fong et al., 2023). The arena was placed inside a chamber enclosed by blackout curtains. RGB-D (RGB + depth) videos were captured at a resolution of 1280X720 pixels and 30 frames per second (fps) under white lighting, using Orbbec Gemini 335L depth camera. To obtain 3-D keypoints, we manually annotated 10 2-D keypoints (see Supporting Information 3) on the RGB frames and projected them to the corresponding 3-D coordinates by aligning these annotations with the reconstructed point-cloud from the depth maps (Fig. 2F). Segmentations were generated using Segment Anything (Kirillov et al., 2023), followed by manual verification and refinement.

**Figure 2.**
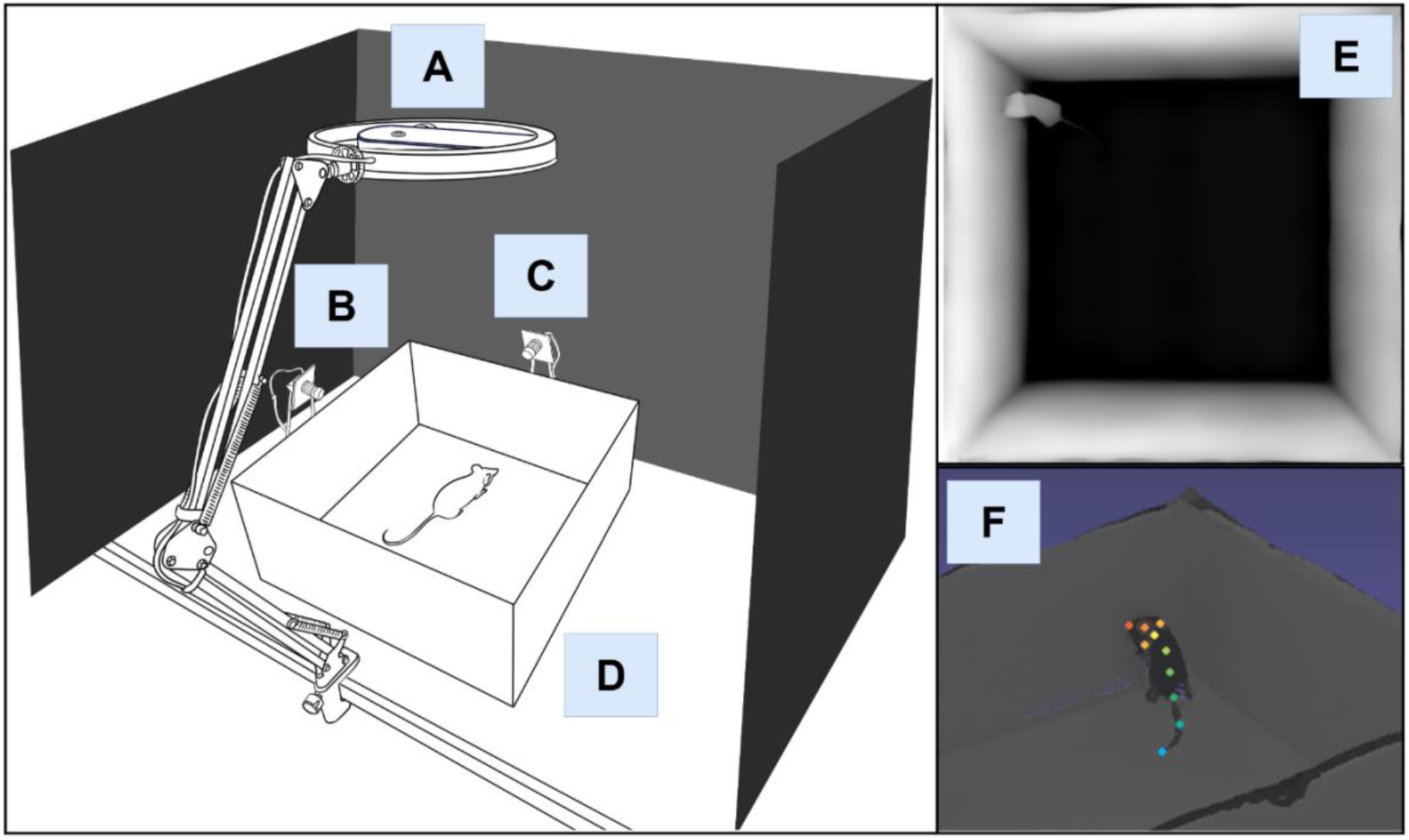
Depth camera setup for recording RGB and depth videos as ground truth verification. (A) Orbbec Gemini 335L structured light depth sensing camera installed at the center of a ring light. (B-C) Optional side webcams, used only to assist in behavior labeling. (D) Open-field arena. (E) An example of the ground-truth depth map after repairing the invalid pixels. Depth values are normalized to 0-255 for better visualization. (F) An example of ground truth 3-D keypoints from projecting the manually labeled 2-D keypoints using the ground truth depth map in E.

To evaluate the generality of our approach we analyzed two additional publicly available resources. First, we re-annotated the MoSeq dataset (Wiltschko et al., 2020), which is comprised of 1.5 h of dual-mouse (FVB/N) recordings captured with a Kinect V1 depth camera at 30 fps in a circular arena. Second, we re-annotated 12, 10-minute videos of single-mouse (C57BL/6) from the PyMouseTracks collection (Fong et al., 2023), acquired under infrared illumination at 960 × 960 pixels and 30 fps with a Raspberry Pi camera. All recordings were labelled with the same 10 keypoints used for our depth-camera dataset. For subsequent behavioral analyses, a representative subset of frames were manually classified—selected via a combination of k-means clustering and interval sampling—into four categories: grooming, rearing, locomotion, and stationary.

### From 2-D Pose & Depth Pseudo Labels to a Posed Mouse Model

To reconstruct 3-D kinematics from 2-D recordings without dedicated depth-capture hardware, we first extracted 2-D keypoints using two commonly used detectors: DeepLabCut-SuperAnimal-Topview (Ye et al., 2024) with resnet-50 backbone, which is optimized for rodent postures, and YOLO11m-pose (Jocher & Qiu, 2024), a lightweight, device-agnostic framework that facilitates cross-platform deployment. Both networks were fine-tuned on a small set of manually annotated frames, and their predictions on full-length videos were post-processed with confidence threshold filtering (0.6).

For the MDE model, we adopted Depth-Anything (Yang et al., 2024), which offers a favorable balance of accuracy and throughput, while also natively supporting metric-scale fine-tuning—essential for quantitative kinematic analysis. Diffusion-based MDE models (e.g., Ke et al. 2024) may recover finer surface details, but their absence of absolute-scale outputs and substantially slower inference currently limit their suitability for large-scale rodent-behavior studies. Crucially, our synthetic-data pipeline is model-agnostic, allowing future advances in monocular depth estimation to be integrated without substantive changes to the workflow.

Because the MDE model was pre-trained on scenes unlike a freely moving mouse in an open-field arena, its raw depth maps exhibit systematic error. We therefore introduced an optimization stage that couples non-uniform rational B-spline (NURBS) spine fitting with biomechanical constraints (see Supporting Information 1 for full derivations). Building on the 3-D mouse model of Bolaños et al. (Bolaños et al., 2021), we developed an automated pipeline with motion-control routine to refine the depth-derived pseudo-labels into anatomically realistic poses.

First, we estimate the overall body scale from the 3-D positions of the ears and the NURBS-fitted spinal curve; the inner-ear distance, which is largely insensitive to adiposity, serves as a stable skeletal reference (Giordano et al., 2022). Next, we apply random “inflation” factors at body-part levels (e.g. buttock) to emulate different levels of body fat. Because limb keypoints are often occluded in top-view videos, plausible limb configurations are synthesized with a set of biomechanically inspired controllers and a data-driven pose sampler trained on side-view footage using the DANNCE dataset (Dunn et al., 2021). A fast mesh-level collision detector then rejects any pose that self-intersects or penetrates the ground. Collectively, these steps convert noisy pseudo labels into anatomically coherent, physically plausible 3-D pose reconstructions even under sparse, occlusion-prone observations. Detailed implementations are presented in our codes and Blender project files.

### Scene Rendering and Style Transfer

To generate high-quality synthetic data across a wide variety of experimental arenas, we created a streamlined scene-modelling workflow in Blender 4.1. Users with computer-aided design (CAD) experience (e.g., those who design 3-D-printed or integrate Thorlabs components) can import pre-designed environments directly as .STL or .STEP files. Entire arenas can also be reconstructed from smartphone videos with photogrammetry tools such as Wonder Studio or Luma AI and imported as textured meshes. After the arena model was loaded, global illumination was set either to a ranged soft-daylight or manually adjusted to match the target recording conditions.

The articulated 3-D mouse model was then placed in the scene, and a virtual top-view camera renders synchronized RGB images, metric depth maps, and semantic-segmentation masks. Key-point annotations were obtained by projecting the joint coordinates of the posed mouse model into image space, guaranteeing pixel-accurate labels. Because fully photorealistic scene construction for every laboratory is impractical, we reduce the synthetic-to-real domain gap by applying an unpaired image-to-image style-transfer network called Unpaired Image Translation via Vector Symbolic Architectures (VSAIT) (Theiss et al., 2022) to the rendered outputs before they are used for training, similar to the previous work (Bolaños et al., 2021). To further reduce such a gap, we also developed new realistic skin/fur textures for flexibly fitting various mouse strains and cranial surgeries.

### Training / Fine-tuning Models

Once the synthetic images with accurate 3-D annotations were available, we fine-tuned the MDE model on this synthetic dataset. To address the specific challenges of rodent recordings—small body size and frequent self-occlusion, we augmented the usual photometric and smoothness terms with a mouse-semantic loss. Conventional depth-estimation losses, developed for human-scale scenes and multi-view indoor datasets, tend to blur small, contiguous structures such as paws, snout, and tail. Our dynamic semantic-weighting scheme down-weights the static arena background and up-weights the small mouse segments, thus preserving fine anatomical detail. Manually assigning pixel-level semantic labels to mouse images—for example, distinguishing every pixel that belongs to the paws, head, and other body parts—is extremely labor-intensive. Accordingly, we leveraged the mesh-to-skeleton correspondence produced by our synthetic pipeline to generate these labels automatically. Our pipeline and loss formulation are readily extensible to applications that demand higher anatomical resolution—such as head-fixed paradigms—because the synthetic renderer can output fine-grained, body-part semantic masks (see Supporting Information 2 for implementation details).

### 3-D Behavioral Classification

Building on the 3-D lifting results, we developed an XGBoost-based (Chen & Guestrin, 2016) pipeline for automated supervised behavior classification and kinematic analysis using xgboost-3.0.0. Behavior classifiers of this type rarely operate on raw key-point time series; instead, they perform best when given defined spatiotemporal kinematic features that summarize posture and motion over time windows (Gharagozloo et al., 2021; Huang et al., 2021; Segalin et al., 2021). Thus, we extracted 74 spatiotemporal features from mouse poses to serve as inputs for a supervised behavior classifier. These PyMouse Lifter features are broadly classified into three categories: motion-, appearance-, and image-based. Motion features quantify locomotion (e.g., speed and directional changes), appearance features characterize the pose within a single frame (e.g., head and body orientation), and image-based features measure pixel-intensity changes between consecutive frames (detailed definitions are provided in Supporting Information 3).

Mouse behavior was manually classified into four ethologically relevant categories—grooming, rearing, locomotion, and stationary. Class imbalance was addressed by enabling xgboost-3.0.0 to rescale the loss function automatically via its scale_pos_weight parameter. Following the one-versus-rest paradigm implemented in MARS (Segalin et al., 2021), we trained one independent binary classifier per category, which allows new behaviors to be added or removed without retraining the entire model suite.

To identify the smallest feature set that supports real-time inference with acceptable accuracy, we quantified feature relevance with two complementary metrics: (i) Impurity-based importance (gain) computed by XGBoost on the combined training + validation folds; (ii) Permutation importance (scikit-learn package) computed on the held-out test set to avoid information leakage. Then we compared models trained on the full complement of spatiotemporal features with models trained on a reduced subset that omitted low-importance variables and long temporal windows. We finally tested whether the streamlined configuration can be feasible for real-time behavior classification on various hardware configurations (Supporting Information 3).

## 3. Results

### PyMouse Lifter and creation of synthetic 3-D training data

Large quantities of rodent behavior have been captured from a single, top-down camera angle—the de-facto standard since Crawley’s protocols (Crawley, 1985)—yet most laboratories still distill these rich videos into coarse read-outs such as arena occupancy or total distance travelled. Unlocking the frame-by-frame animal 3-D kinematics hidden within this footage, and doing so with latencies compatible with closed-loop perturbation, would markedly accelerate circuit-level studies of behavior. Current neural-network based solutions promise such reconstructions with a single top-view camera (Gosztolai et al., 2021; Hu et al., 2023), but only after researchers collect dedicated multi-view or depth-camera training datasets—a costly and inflexible prerequisite that can be difficult to adopt.

To address this need, we developed a 2-D-to-3-D data generation pipeline that employs synthetic data augmentation and leverages a mouse body model (Fig. 1; details in Methods) and data from an overhead camera (Fig. 2) without the need to pre-calibrate the arena. Beginning with a small set of manually annotated frames, a trained 2-D pose estimation model produced coarse 2-D keypoints (Fig. 3B) for unlabeled images (Fig. 3A). The approximate 3-D keypoints were reconstructed (Fig. 3C) by combining 2D keypoints with coarse depth maps (Fig. 3B) derived from a pretrained MDE model (Depth Anything). We then refined the skeletons of our mouse model with an automated motion-control system that enforces biomechanical constraints, driving an anatomically accurate mouse mesh to specific poses using inverse kinematics (Fig. 3D; details in Supporting Information 1). Although the poses are not pixel-exact replicas of the source frames, their anatomical fidelity is based on a model of the mouse body and sufficient to render synthetic 3D data for retraining the neural network models (MDE and 2D pose estimators).

**Figure 3.**
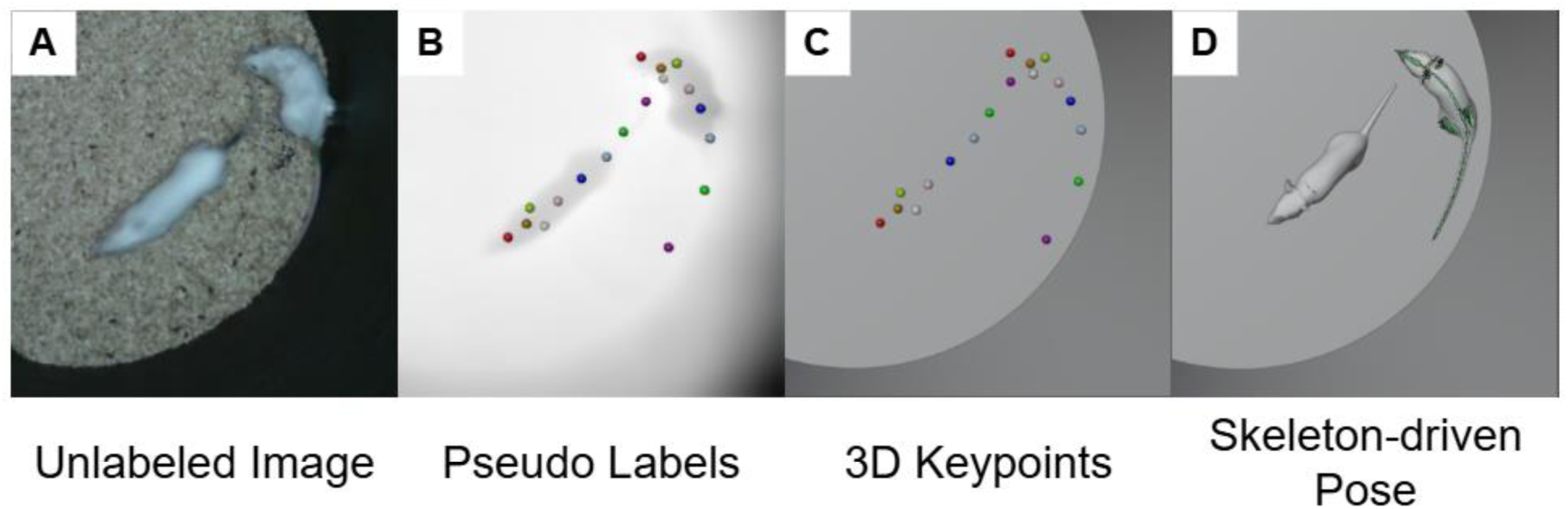
Detailed workflow for generating 3-D pose from real 2-D images, illustrated using MoSeq dataset (Wiltschko et al., 2020). (A) Starting with an unlabeled real image. (B) 2-D keypoints from the YOLO11m-pose estimation model (trained on a small number of manually labeled samples) and a coarse depth map from a pre-trained Depth Anything model. (C) Coarse 3-D keypoints in the synthetic scene. (D) Optimized skeleton-driven pose using the information from estimated 3-D keypoints and biomechanical constraints.

Once realistic synthetic mouse poses were generated, we employed computer graphics (CG) rendering combined with style transfer techniques to produce high-fidelity synthetic data. This approach significantly reduced the domain gap between the training data and real-world conditions (Fig. 4). Specifically, the 3-D mouse model was placed in a virtual scene and customized with strain-specific traits—such as eye morphology and fur texture—to closely match the target mouse phenotype (see Methods).

**Figure 4.**
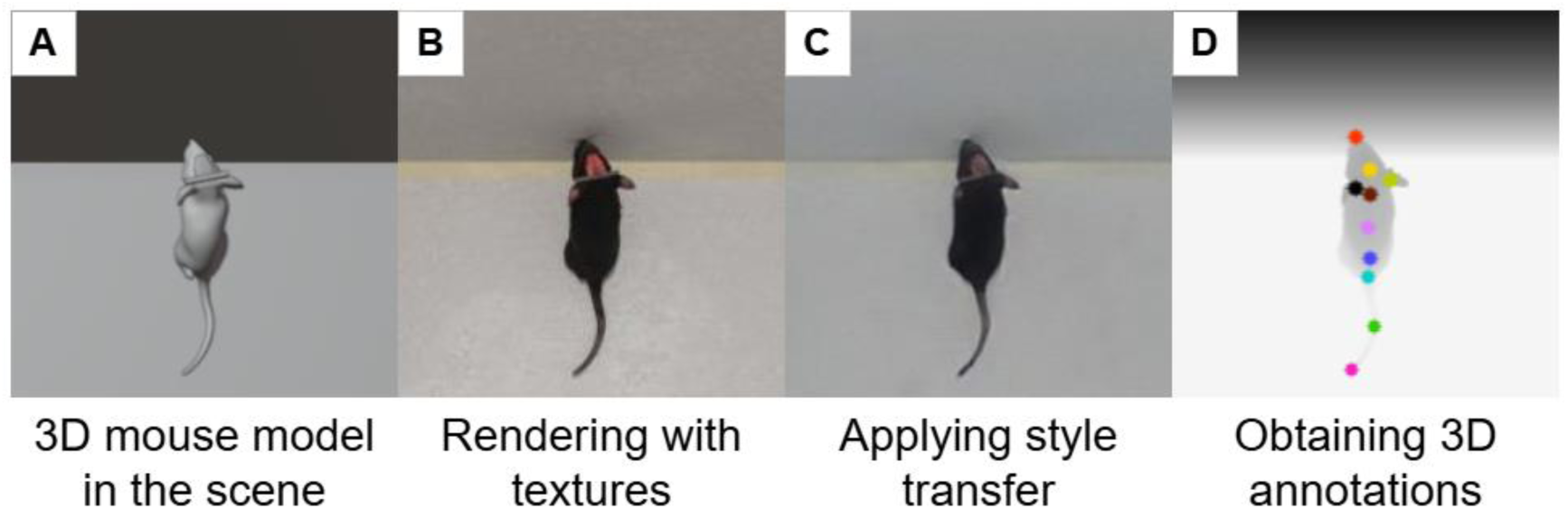
Detailed workflow and example for synthesizing labeled training images from a posed 3-D mouse model, shown with our single-mouse open-field dataset (cranial-window stroke mouse). (A) Posed 3-D mouse model generated by automated scripting. (B) Rendering with realistic skin- and fur-like materials matching the target mouse. (C) Style-transfer refinement of the renders. (D) Dense, unbiased annotations—keypoints and depth maps—exported from the scene.

The resulting synthetic images, along with their error-free annotations, served as the primary training dataset for our fine-tuned depth estimation model. Notably, we did not directly use pseudo-labeled raw images (e.g., as in *Depth Anything* [Li et al., 2024]) for training. Instead, we leveraged 3-D synthetic mouse priors to rationalize the pseudo labels. Although the corrected 3-D keypoints were kinematically and anatomically plausible, they exhibited an average error of 24.38 ± 18.56 mm (mean ± SD, same for all below) compared to the ground-truth 3-D keypoints in the original images (Supporting Information 7) and should be thought of as a vehicle for model training and not an accurate 3D representation of original mouse. Overall, we generated high-quality 3-D training data entirely from synthetic images (based on lifted 2-D information from real data), eliminating the need for specialized 3-D-capture hardware to collect 3-D training datasets.

We measured the ability of the synthetic training data to support the pose and MDE models using single- and dual-animal real test datasets. To evaluate the impact of style-transfer post-processing, we compared model performance on real-world data with and without applying this step to the synthetic images for training (Fig. 5). Specifically, we compared 4 model variants: (i) a zero-shot baseline using an off-the-shelf MDE model with no rodent-specific retraining, (ii) a model fine-tuned on real images, (iii) a model fine-tuned on unaltered rendered images, and (iv) a model fine-tuned on style-transferred renders.

**Figure 5.**
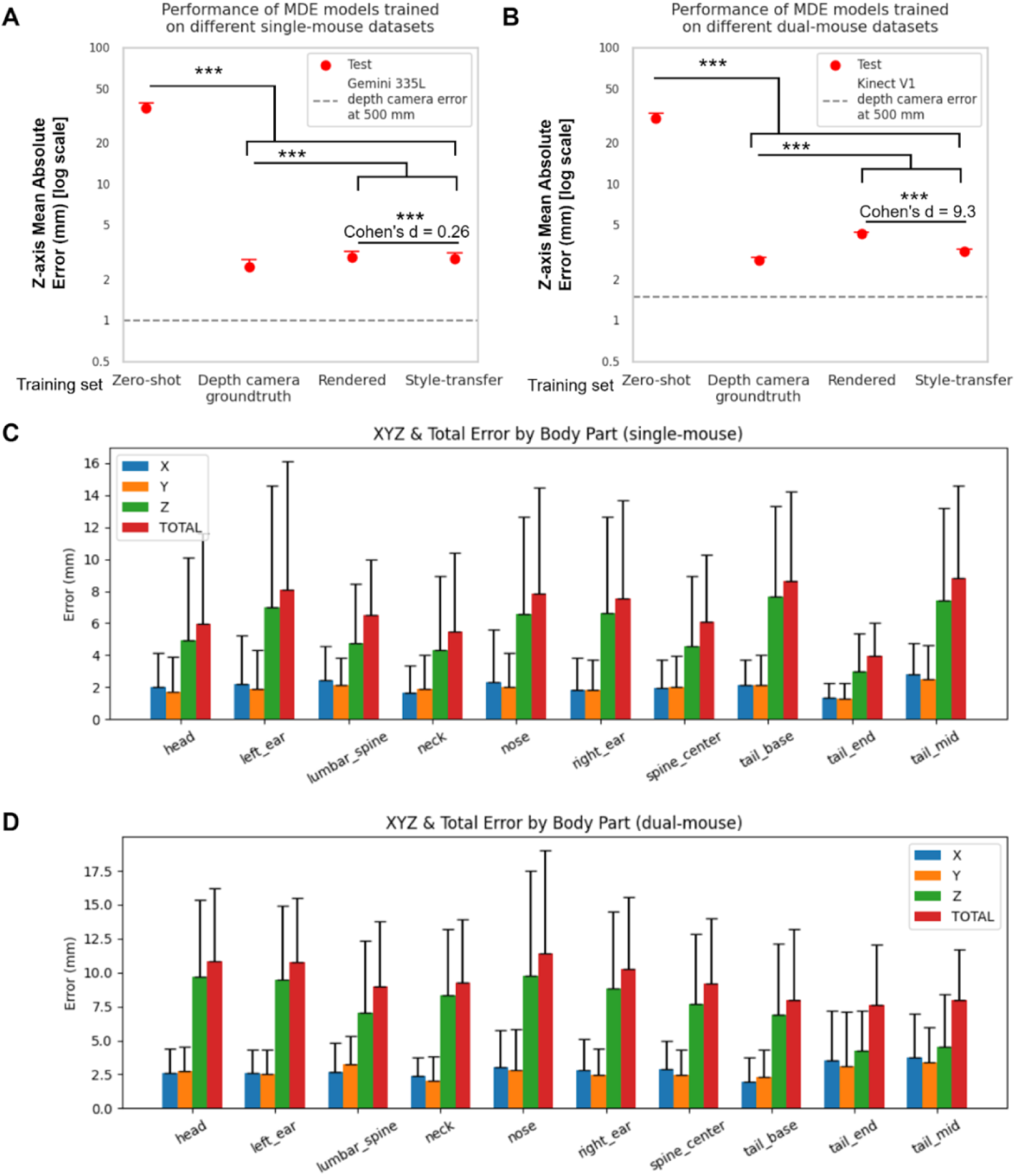
Comparison of MDE models trained on real / rendered synthetic data / style-transferred synthetic data on actual depth camera measurements, and the 2D-to-3D lifting results based on the best of them using the style-transfer module. Test MAEs were calculated as the difference in Z axis from the depth maps taken by the structured light depth camera. The predicted depth map is aligned with the actual depth map at the ground level, and only the valid pixels belonging to the mouse are identified and used for calculation. Depth camera error baselines were obtained from the device datasheets. Stars indicate significant paired t-tests results. The error bars are ± 1 SD. (A) MAEs on our single-mouse openfield dataset (22 mice, each with 1-3 10-min videos; upper error bar shown). (B) MAEs on MoSeq (Wiltschko et al., 2020) dual-mice openfield dataset (2 mice in a 1.5-hour video). (C) 2D-to- 3D projection errors on our single-mouse openfield dataset. (D) 2D-to-3D projection errors on MoSeq dual-mice openfield dataset.

Because existing pretrained MDE models are optimized for human-centered computer-vision benchmarks, they provide little prior knowledge of rodent morphology; the zero-shot baseline therefore serves as a lower bound for accuracy, illustrating how our model-based synthetic pipeline transforms noisy pseudo-labels into accurate depth predictions.

Fine-tuning the MDE model on synthetic images produced by PyMouse Lifter markedly decreased reconstruction error—from > 30 mm in the zero-shot baseline to ≈ 2.9 mm after fine-tuning—and yielded parallel gains on all auxiliary metrics (Fig. 5A, B; Supporting Information 4). For our single-animal open-field recordings (720X720), the photorealism of the rendered images was already sufficient to obtain near-maximal accuracy when the retrained model was evaluated against ground-truth depth maps (Fig. 5A; Supporting Information 4); consequently, an additional style-transfer stage, although itself a form of data augmentation, conferred no measurable benefit on this dataset. On 1250 test images, the mean difference of the Z-axis mean absolute error (MAE) between the two models was 0.08 mm, which was statistically significant (p < 0.001), with a 95% CI of 0. However, the effect size Cohen’s d = 0.26 is small, and the magnitude of the mean difference is less than 10% of the depth camera hardware error (Fig. 5A).

In contrast, using the MoSeq dataset (Wiltschko et al., 2020)—captured with legacy Kinect V1 depth camera at lower resolution (640X480) and against more visually cluttered backgrounds, we did observe significant benefit from style-enhanced synthetic data: fine-tuning the model using the style-transferred images further reduced mean absolute depth error and improved all evaluation scores (Fig. 5B; Supporting Information 4). Across the 1250-image test set, adding the style-transfer module lowered the Z-axis MAE from 4.31 ± 0.13 mm to 3.19 ± 0.11 mm —a 1.12 mm (-26%) reduction that was highly significant (p < 0.001) and corresponded to an extremely large effect size (Cohen’s d = 9.3). These results indicate that the style-transfer module may shorten the gap between the synthetic images and the real images more when the recording source footage is noisy or the raw rendering style is not photorealistic enough. More details, including standard computer vision metrics for MDE, are presented in Supporting Information 4.

After projecting the 2D keypoints to 3D space, we observed overall 3-D errors of 6.82 ± 5.62 mm for the single-animal open field dataset (Fig. 5C), and 9.43 ± 5.27 mm for the MoSeq dual-animal dataset (Fig. 5D) with YOLO11m-pose model. Although the projection performance of the DeepLabCut model was comparable (Supporting Information 8), we chose to develop our real-time behavioral classification pipeline based on YOLO due to improved python environmental compatibility with Depth Anything. The mean error for each keypoint was under 10 mm. Considering that the average body length of adult laboratory mice (≥12 weeks) is approximately 9–10 cm (Chakraborty et al., 2017; Kidwell et al., 1979), the relative error remains below 10%. These numbers suggest room for improvement, as the 2D-to-3D projection process amplifies errors along the Z-axis. This is partly because MDE models can achieve relatively low error when pixel correspondences are perfectly known; however, when projecting estimated 2D keypoints onto 3D space using predicted depth maps, the uncertainty in keypoint localization compounds the depth estimation error, leading to greater inaccuracies specifically along the Z dimension (explaining differences between values presented in Fig. 5 A,B versus Fig. 5 C,D). Despite these issues, our next-step results (Fig. 6 below) demonstrate that the accuracy is sufficient for downstream behavioral classification tasks.

**Figure 6.**
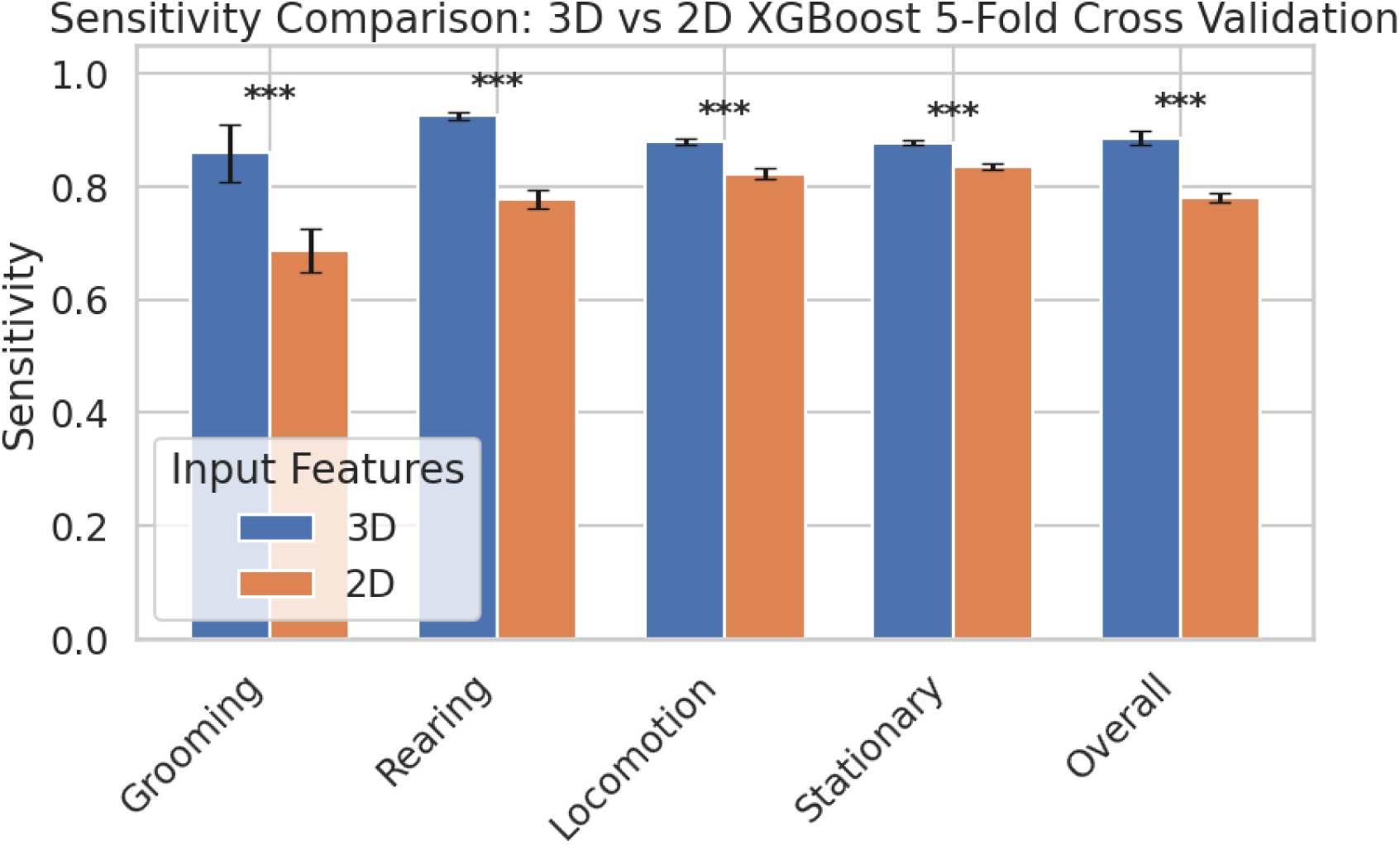
Bar graph of the classification sensitivities comparison between XGBoost models using 3D features or 2D features (see Methods and Supplementary Information 3). The error bars are ± standard deviation.

### Supervised behavior classification with 3-D features

Three-dimensional key-point estimation was motivated by the premise that depth cues encode behaviorally relevant information that 2-D tracking cannot capture. To determine whether this additional z-dimension improves behavioral read-outs, we trained two otherwise identical XGBoost pipelines. The 3-D classifier utilized the full set of 74 defined spatiotemporal features (e.g. nose height, spinal curviness, etc.) extracted from the lifted 3-D keypoints as input, whereas the 2-D baseline was limited to the 49 features that remain after all z-dependent terms were removed (see Methods and Supplementary Information 3). Both models followed a one-versus-rest strategy to classify grooming, rearing, locomotion, and stationary states.

During the labeling stage, we employed interval sampling on the time series data. The labeled frames were initially partitioned into non-overlapping temporal windows to minimize correlations between adjacent frames, thereby reducing the risk of highly similar data appearing in both the training and test sets. We implemented a nested 5-fold cross-validation approach (Cawley & Talbot, 2010) at the window level. In each iteration, one fold was reserved exclusively for testing, while the remaining four folds were allocated for model training (75%) and validation (25%). Performance metrics—including sensitivity and the F1-score—were computed and subsequently averaged across all held-out folds. The results are summarized in Figure 6 and Supporting Information 6.

The 3-D model achieved significantly higher accuracy across all four behaviors, with especially pronounced gains for grooming and rearing, whose kinematics involve substantial vertical movement (Fig. 6). Even under the highly imbalanced class distribution dominated by locomotion and stationary (Supporting Information 5), the sensitivities and F1-scores for grooming and rearing improved significantly (p < 0.001, paired t-tests), confirming that 3-D lifting provides critical information absent from 2D top-down video alone.

To identify which lifted features drove these improvements, we computed feature importance for the multi-class 3-D model using both gain-based and permutation methods (Fig. 7A,B). Despite minor differences between the two estimates, several kinematic descriptors that explicitly depend on depth—such as nose/head height (nose_3d_z / head_3d_z) and body-centric pitch (spine_center_z)—consistently ranked among the top contributors, underscoring the behavioral relevance of the recovered z-axis.

**Figure 7.**
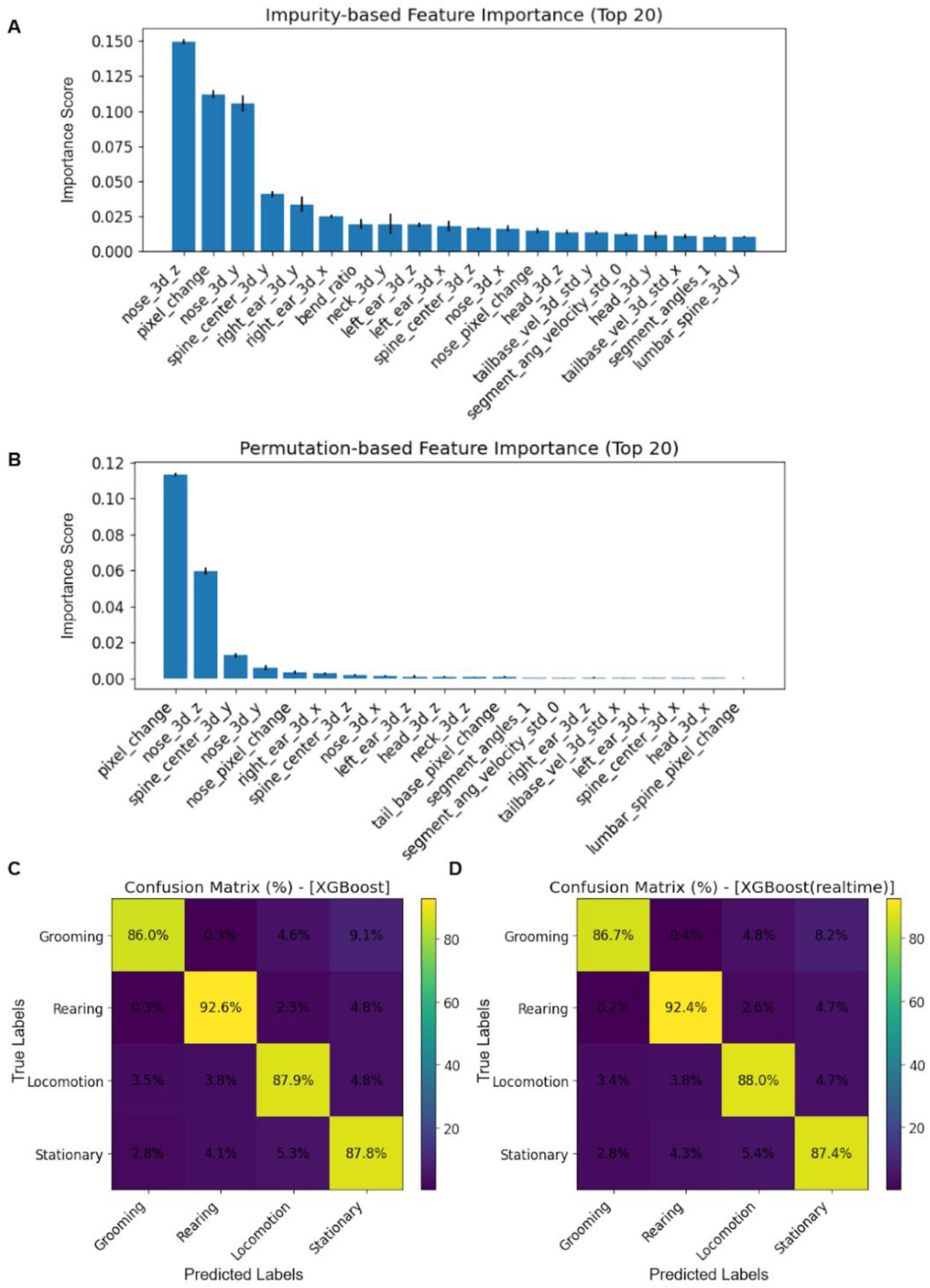
Performance of the assembled multi-class 3-D lifting XGBoost model and ability to do real time classification. (A) Average impurity-based feature importance; (B) Average permutation-based feature importance; (C) 5-fold cross validation confusion matrix; (D) 5-fold cross validation confusion matrix of the lightened assembled multi-class 3-D lifting XGBoost model for real-time processing, using only the 29 critical features from the two adjacent frames (about 33 ms interval at 30 fps).

### Realtime behavior classification using fewer temporal features

To evaluate whether our pipeline can support real-time, closed-loop applications, we retrained a streamlined XGBoost model that retained only the 29 most informative features—those originating from the current and immediately preceding frames and ranking highest in both impurity-based and permutation importance analyses. Eliminating long time-window descriptors and low-contribution variables reduced the feature count by >60 % without sacrificing predictive power: five-fold cross-validation revealed virtually unchanged performance across all behaviors and a modest improvement in grooming detection (Fig. 7C, D). We achieved end-to-end behavioral classification at above 30 fps from raw 2-D images, through 2-D keypoint extraction, depth estimation lifting, and XGBoost across multiple hardware configurations (Sup. Table 3). These results demonstrate that the proposed feature set and model architecture are compatible with real-time deployment.

## 4. Discussion

### Conclusion

Deep-learning-based automated pose estimation systems are increasingly utilized by biomedical researchers and have been demonstrated to be applicable in a variety of behavioral experiments (Dunn et al., 2021; Mathis et al., 2018; Pereira et al., 2022; Weinreb et al., 2024). However, a major limitation of data-driven deep learning frameworks is that they require a substantial amount of annotated training data to achieve efficient automation. Although significant advances have been made in pre-training and zero-shot estimation in the field of computer vision (Yang et al. 2024; Yan et al. 2021; Ke et al. 2024), large-scale, high-quality datasets necessary for developing large models are generally lacking in fundamental research involving experimental animals, which is not driven by industrial or commercial forces.

This work demonstrates that accurate 3-D pose estimation and behavior classification can be achieved with nothing more than a monocular webcam and sparse 2-D key-point labels. By combining a lightweight generative synthesis pipeline with lifted 3-D key-points, we overcame the usual bottlenecks of specialized hardware, dense annotations, and large training corpora that have limited the deployment of 3-D methods in animal open-field assays. Moreover, since our method imposes no strict requirements on data acquisition, it can also be retroactively applied to previously collected or archival 2-D video data, enabling enriched reanalysis and deeper insight without the need for additional experiments. The extra depth cues translate into measurable gains in downstream behavior decoding and, in principle, enable real-time closed-loop manipulations of select behaviors in freely moving mice. Previously, other work has reported 3-D keypoint estimation from a single camera, but this required users to set up calibrated 3-D capture devices to acquire initial training data (Hu et al., 2023). We suggest that the data augmentation in our pipeline and the use of a 3-D mouse model (in essence a synthetic 3-D ground truth) helps to overcome dependence on a calibrated 3D arena. The values of 3-D errors were lower (and improved) over the visible keypoint errors reported in Hu et al. (2023). However, because our keypoint definitions are based on Weinreb et al. (Weinreb et al., 2024) and differ substantially from those used by Hu et al. (2023), a direct comparison is limited.

Our approach also sidesteps several practical constraints that accompany structured-light depth cameras. To capture RGB-D image streams at sufficient speed to prevent motion blur and avoid missing fast movements, it is often necessary to restrict the disparity search range. This constraint improves computational efficiency and reduces the likelihood of mismatches. However, it also introduces trade-offs in depth resolution. For example, devices such as the Orbbec Gemini and Intel RealSense series typically require a minimum effective imaging distance of approximately 0.5 meters when operating with a 128-level disparity search range (Orbbec, 2025; Intel, 2023) for stable 30fps recording. Pushing to 60 fps is possible with the cost of 64-level disparity search range that increases the minimum working distance to 1 meter, which sacrifices the details of the mouse and may not work in homecages where space is limited. Moreover, shiny bedding, acrylic walls, and metal implants scatter the projected pattern, generating holes and spikes in the depth map and often preventing the use of a protective cage lid. In contrast, our pipeline infers depth directly from monocular RGB video. It therefore eliminates IR-pattern line-of-sight issues, operates at the full frame rate of any commodity camera, and scales from cramped arenas to open fields without recalibration or specialized lighting, all while preserving video-rate throughput.

### Limitations and future directions

Our current 2-D-to-3-D synthesis pipeline is confined to top-view recordings of freely moving mice. Although we prototyped a bottom-view variant—where limb kinematics are easier to drive from paw key-points—the approach was hampered by the poor generalization of off-the-shelf MDE models (Ranftl et al., 2022; Bhat et al., 2023; Yang et al., 2024). The domain gap between typical training data (human-centered, outdoor scenes) and mouse ventral images appears even larger than for dorsal views, so the bottom view model would require a sizable, manually curated seed dataset before synthetic augmentation could be effective. We encountered comparable difficulties in head-fixed grasping experiments, where occlusions and reflective surfaces further degraded monocular depth accuracy.

More broadly, our workflow has so far been trained and evaluated in the controlled visual context of open-field arenas. Extending it to richer environments (complex mazes, multi-animal settings, variable lighting) will likely demand (i) richer geometric cues—e.g., fused RGB-D point clouds or multi-view stereo—and (ii) larger collections of real videos to co-evolve with the synthetic generator. We envisage an iterative cycle in which new real footage guides simulation fidelity, while progressively refined synthetic data reduces annotation burden.

Despite these constraints, we believe that our open-source 3-D synthetic mouse model, alongside the provided codebase and annotated dataset, offers a valuable and practical starting point for the broader behavioral neuroscience and computer vision communities. While this study primarily employed a supervised learning framework to intuitively quantify the behavioral analysis gains of 3-D over 2-D data, we acknowledge that unsupervised approaches such as keypoint-MoSeq (Weinreb et al., 2024) and B-SOiD (Hsu & Yttri, 2021) may also benefit significantly from our pipeline. These methods could leverage our 3-D representations to discover latent behavioral motifs with higher fidelity and potentially greater interpretability (Weinreb et al., 2024). By lowering the barrier to high-precision pose annotation, it should accelerate both methodological advances (e.g., point-cloud-based pose estimation, self-supervised behavioral discovery) and biological discoveries in rodent behavioral study.

## Acknowledgments

This work was supported by a Canadian Institutes of Health Research (CIHR) Foundation Grant FDN-143209 and project grant PJT-180631 to T.H.M. THM was also supported by the Brain Canada Neurophotonics Platform, a Natural Science and Engineering Council of Canada (NSERC; GPIN-2022-03723). This work was supported by resources made available through the Dynamic Brain Circuits cluster and the NeuroImaging and NeuroComputation Centre at the UBC Djavad Mowafaghian Centre for Brain Health (RRID SCR_019086) and made use of the DataBinge forum. We are especially grateful to Dr. Sandeep R. Datta and his laboratory at Harvard Medical School for generously providing access to behavioral datasets from their MoSeq (2020) and keypoint-MoSeq (2024) studies.

## 6. Data Availability

All code and models will be available on public repositories: https://github.com/Haozong-Zeng/PyMouse-Lifter and [video data storage] once published; see workflow document for an overview of software steps. Before publication only the DEMOs are released.

## Bibliography

Bhat, S. F., Birkl, R., Wofk, D., Wonka, P., & Müller, M. (2023). ZoeDepth: Zero-shot transfer by combining relative and metric depth. In arXiv [cs.CV]. http://arxiv.org/abs/2302.12288

Bolaños, L. A., Xiao, D., Ford, N. L., LeDue, J. M., Gupta, P. K., Doebeli, C., Hu, H., Rhodin, H., & Murphy, T. H. (2021). A three-dimensional virtual mouse generates synthetic training data for behavioral analysis. Nat. Methods, 18(4), 378–381. 10.1038/s41592-021-01103-9

Cawley, G., & Talbot, N. L. C. (2010). On over-fitting in model selection and subsequent selection bias in performance evaluation. J. Mach. Learn. Res., 11, 2079–2107. 10.5555/1756006.1859921

Chakraborty, R., Park, H. N., Tan, C. C., Weiss, P., Prunty, M. C., & Pardue, M. T. (2017). Association of body length with ocular parameters in mice. Optom. Vis. Sci., 94(3), 387–394. 10.1097/OPX.0000000000001036

Chen, T., & Guestrin, C. (2016, August 13). XGBoost: A Scalable Tree Boosting System. Proceedings of the 22nd ACM SIGKDD International Conference on Knowledge Discovery and Data Mining, New York, NY, USA. 10.1145/2939672.2939785

Crawley, J. N. (1985). Exploratory behavior models of anxiety in mice. Neurosci. Biobehav. Rev., 9(1), 37–44. 10.1016/0149-7634(85)90030-2

Dunn, T. W., Marshall, J. D., Severson, K. S., Aldarondo, D. E., Hildebrand, D. G. C., Chettih, S. N., Wang, W. L., Gellis, A. J., Carlson, D. E., Aronov, D., Freiwald, W. A., Wang, F., & Ölveczky, B. P. (2021). Geometric deep learning enables 3D kinematic profiling across species and environments. Nat. Methods, 18(5), 564–573. 10.1038/s41592-021-01106-6

Fong, T., Hu, H., Gupta, P., Jury, B., & Murphy, T. H. (2023). PyMouseTracks: Flexible computer vision and RFID-based system for multiple mouse tracking and behavioral assessment. eNeuro, 10(5). 10.1523/ENEURO.0127-22.2023

Gharagozloo, M., Amrani, A., Wittingstall, K., Hamilton-Wright, A., & Gris, D. (2021). Machine learning in modeling of mouse behavior. Front. Neurosci., 15, 700253. 10.3389/fnins.2021.700253

Giordano, A., Cinti, F., Canese, R., Carpinelli, G., Colleluori, G., Di Vincenzo, A., Palombelli, G., Severi, I., Moretti, M., Redaelli, C., Partridge, J., Zingaretti, M. C., Agostini, A., Sternardi, F., Giovagnoni, A., Castorina, S., & Cinti, S. (2022). The adipose organ is a unitary structure in mice and humans. Biomedicines, 10(9), 2275. 10.3390/biomedicines10092275

Gosztolai, A., Günel, S., Lobato-Ríos, V., Pietro Abrate, M., Morales, D., Rhodin, H., Fua, P., & Ramdya, P. (2021). LiftPose3D, a deep learning-based approach for transforming two-dimensional to three-dimensional poses in laboratory animals. Nat. Methods, 18(8), 975–981. 10.1038/s41592-021-01226-z

Hatton-Jones, K. M., Christie, C., Griffith, T. A., Smith, A. G., Naghipour, S., Robertson, K., Russell, J. S., Peart, J. N., Headrick, J. P., Cox, A. J., & du Toit, E. F. (2021). A YOLO based software for automated detection and analysis of rodent behaviour in the open field arena. Comput. Biol. Med., 134(104474), 104474. 10.1016/j.compbiomed.2021.104474

Hsu, A. I., & Yttri, E. A. (2021). B-SOiD, an open-source unsupervised algorithm for identification and fast prediction of behaviors. Nat. Commun., 12(1), 5188. 10.1038/s41467-021-25420-x

Hu, B., Seybold, B., Yang, S., Sud, A., Liu, Y., Barron, K., Cha, P., Cosino, M., Karlsson, E., Kite, J., Kolumam, G., Preciado, J., Zavala-Solorio, J., Zhang, C., Zhang, X., Voorbach, M., Tovcimak, A. E., Ruby, J. G., & Ross, D. A. (2023). 3D mouse pose from single-view video and a new dataset. Sci. Rep., 13(1), 13554. 10.1038/s41598-023-40738-w

Huang, R., Nikooyan, A. A., Xu, B., Joseph, M. S., Damavandi, H. G., von Trotha, N., Li, L., Bhattarai, A., Zadeh, D., Seo, Y., Liu, X., Truong, P. A., Koo, E. H., Leiter, J. C., & Lu, D. C. (2021). Machine learning classifies predictive kinematic features in a mouse model of neurodegeneration. Scientific Reports, 11(1), 1–16. 10.1038/s41598-021-82694-3

Karashchuk, P., Rupp, K. L., Dickinson, E. S., Walling-Bell, S., Sanders, E., Azim, E., Brunton, B. W., & Tuthill, J. C. (2021). Anipose: A toolkit for robust markerless 3D pose estimation. Cell Rep., 36(13), 109730. 10.1016/j.celrep.2021.109730

Kidwell, J. F., Herbert, J. G., & Chase, H. B. (1979). The inheritance of growth and form in the mouse. V. Allometric growth. Growth, 43(1), 47–57.

Mathis, A., Mamidanna, P., Cury, K. M., Abe, T., Murthy, V. N., Mathis, M. W., & Bethge, M. (2018). DeepLabCut: Markerless pose estimation of user-defined body parts with deep learning. Nat Neurosci, 21(9), 1281–1289. 10.1038/s41593-018-0209-y

Pereira, T. D., Tabris, N., Matsliah, A., Turner, D. M., Li, J., Ravindranath, S., Papadoyannis, E. S., Normand, E., Deutsch, D. S., Wang, Z. Y., McKenzie-Smith, G. C., Mitelut, C. C., Castro, M. D., D’Uva, J., Kislin, M., Sanes, D. H., Kocher, S. D., Wang, S. S.-H., Falkner, A. L., … Murthy, M. (2022). SLEAP: A deep learning system for multi-animal pose tracking. Nat. Methods, 19(4), 486–495. 10.1038/s41592-022-01426-1

Ranftl, R., Lasinger, K., Hafner, D., Schindler, K., & Koltun, V. (2022). Towards robust monocular depth estimation: Mixing datasets for zero-shot cross-dataset transfer. IEEE Trans. Pattern Anal. Mach. Intell., 44(3), 1623–1637. 10.1109/TPAMI.2020.3019967

Segalin, C., Williams, J., Karigo, T., Hui, M., Zelikowsky, M., Sun, J. J., Perona, P., Anderson, D. J., & Kennedy, A. (2021). The Mouse Action Recognition System (MARS) software pipeline for automated analysis of social behaviors in mice. Elife, 10. 10.7554/eLife.63720

Sun, G., Lyu, C., Cai, R., Yu, C., Sun, H., Schriver, K. E., Gao, L., & Li, X. (2021). DeepBhvTracking: A novel behavior tracking method for laboratory animals based on deep learning. Front. Behav. Neurosci., 15, 750894. 10.3389/fnbeh.2021.750894

Weinreb, C., Pearl, J. E., Lin, S., Osman, M. A. M., Zhang, L., Annapragada, S., Conlin, E., Hoffmann, R., Makowska, S., Gillis, W. F., Jay, M., Ye, S., Mathis, A., Mathis, M. W., Pereira, T., Linderman, S. W., & Datta, S. R. (2024). Keypoint-MoSeq: Parsing behavior by linking point tracking to pose dynamics. Nat. Methods, 21(7), 1329–1339. 10.1038/s41592-024-02318-2

Wiltschko, A. B., Tsukahara, T., Zeine, A., Anyoha, R., Gillis, W. F., Markowitz, J. E., Peterson, R. E., Katon, J., Johnson, M. J., & Datta, S. R. (2020). Revealing the structure of pharmacobehavioral space through motion sequencing. Nat Neurosci, 23(11), 1433– 1443. 10.1038/s41593-020-00706-3

Ye, S., Filippova, A., Lauer, J., Schneider, S., Vidal, M., Qiu, T., Mathis, A., & Mathis, M. W. (2024). SuperAnimal pretrained pose estimation models for behavioral analysis. Nat. Commun., 15(1), 5165. 10.1038/s41467-024-48792-2

Zuffi, S., Kanazawa, A., Jacobs, D., & Black, M. J. (2017). 3-D menagerie: Modeling the 3-D shape and pose of animals. Proceedings of the IEEE Conference on Computer Vision and Pattern Recognition (CVPR). 10.48550/arXiv.1611.07700

Loper, M., Mahmood, N., Romero, J., Pons-Moll, G., & Black, M. J. (2015). SMPL: A skinned multi-person linear model. ACM Transactions on Graphics (TOG), 34(6), Article 248, 1–16. 10.1145/2816795.2818013

Jocher, G., & Qiu, J. (2024). Ultralytics YOLO11 (Version 11.0.0) [Computer software]. Ultralytics. https://github.com/ultralytics/ultralytics

Yang, L., Kang, B., Huang, Z., Xu, X., Feng, J., & Zhao, H. (2024). Depth Anything: Unleashing the power of large-scale unlabeled data. In Proceedings of the IEEE/CVF Conference on Computer Vision and Pattern Recognition (CVPR). arXiv preprint arXiv:2401.10891. 10.48550/arXiv.2401.10891

Yang, L., Kang, B., Huang, Z., Zhao, Z., Xu, X., Feng, J., & Zhao, H. (2024). Depth Anything V2. In Proceedings of the 38th Conference on Neural Information Processing Systems (NeurIPS).

Yan, H., Chen, J., Zhang, X., Zhang, S., Jiao, N., Liang, X., & Zheng, T. (2021). UltraPose: Synthesizing dense pose with 1 billion points by human-body decoupling 3-D model. In Proceedings of the IEEE/CVF International Conference on Computer Vision (ICCV). arXiv preprint arXiv:2110.15267. 10.48550/arXiv.2110.15267

Li, S., Günel, S., Ostrek, M., Ramdya, P., Fua, P., & Rhodin, H. (2020). Deformation-aware unpaired image translation for pose estimation on laboratory animals. In Proceedings of the IEEE/CVF Conference on Computer Vision and Pattern Recognition (CVPR). arXiv preprint arXiv:2001.08601. 10.48550/arXiv.2001.08601

Ke, B., Obukhov, A., Huang, S., Metzger, N., Caye Daudt, R., & Schindler, K. (2024). Repurposing diffusion-based image generators for monocular depth estimation. In Proceedings of the IEEE/CVF Conference on Computer Vision and Pattern Recognition (CVPR). arXiv preprint arXiv:2312.02145. 10.48550/arXiv.2312.02145

Orbbec. (n.d.). Gemini 330 Series datasheet [Datasheet]. Retrieved July 21, 2025, from https://www.orbbec.com/docs/g330-orbbec-gemini-330-series-datasheet/, 117–125. 10.1038/s41592-018-0234-5

Intel. (2023, March). Intel RealSense D400 Series Datasheet [Datasheet]. Retrieved July 21, 2025, from https://www.intel.com/content/www/us/en/products/sku/190004/intel-realsense-depth-camera-d435i/specifications.html

