## Supporting Information 1-8 for "PyMouse Lifter: Real Time 3-D Pose Estimation for Mice with Only 2-D Annotation Via Data Synthesis"

**Supporting Information 1: Synthetic Mouse Optimization**

At a certain frame t of the video, the 3-D key points $\mathbf{K}_{\text{spines}}$ of the mouse spine are:

$$\mathbf{K}_{\text{spine,t}} = [\mathbf{k}_{\text{nose,t}}, \mathbf{k}_{\text{head,t}}, \dots, \mathbf{k}_{\text{tail,t}}]$$

Let n denote the number of keypoints on the head-to-tail spine. The approximate length of the spine curve $\mathrm{L_t}$ can be obtained by accumulating the Euclidean distance between adjacent key points:

$$L_t = \sum_{i=1}^{n-1} \| \boldsymbol{k}_{i+1} - \boldsymbol{k}_i \| = \sum_{i=1}^{n-1} \sqrt{(x_{i+1} - x_i)^2 + (y_{i+1} - y_i)^2 + (z_{i+1} - z_i)^2}$$

To align the Z-axis coordinates zi and minimize the variation in spinal curve length $\mathrm{L_t}$ across frames, we apply a unified linear transformation:

$$z_i' = a \cdot z_i + b$$

where: a is a scaling factor, and b is a translation factor.

The adjusted spinal curve length after transformation becomes:

$$L_t' = \sum_{i=1}^{n-1} \sqrt{(x_{i+1} - x_i)^2 + (y_{i+1} - y_i)^2 + (z_{i+1}' - z_i')^2}$$

The goal is to minimize the total variation in spinal curve lengths across all frames in the video:

$$\text{Minimize} \sum_{t=1}^{T} | L_t' - L_0 |$$

where $L_0$ is the reference spinal curve length, ideally kept constant across frames. For simplicity we use the average spinal curve length across all frames:

$$L_0 = \frac{1}{T} \sum_{t=1}^{T} L_t$$

I then use gradient descent to minimize the objective function and apply the optimized linear transformation to the Z-axis coordinates. Subsequently, a non-uniform rational B-spline (NURBS) curve is fitted to model spinal dynamics.

Given the aligned keypoints of the spine $\mathbf{K}_{\text{spine}} = [\mathbf{k}_{\text{nose}}, \mathbf{k}_{\text{head}}, \dots, \mathbf{k}_{\text{tail}}]$, we fit a NURBS curve $C(u)$ to model the dynamics of the spine:

$$C(u) = \frac{\sum_{i=0}^{n} N_{i,p}(u) \cdot w_i \cdot P_i}{\sum_{i=0}^{n} N_{i,p}(u) \cdot w_i}$$

where:

$N_{i,p}(u)$ is the B-spline basis function of degree p,

$w_i$ is the weight of control point $\boldsymbol{k}_i$.

This step further reduces frame-to-frame errors introduced by the MDE model through curve smoothing. The pseudo-labeled 3-D keypoints of the ears are utilized to guide the orientation of the head.

Given the 3-D positions of the left and right ear keypoints $\mathbf{k}_{\text{left\_ear}}$ and $\mathbf{k}_{\text{right\_ear}}$, the ear vectors are computed as:

$$\mathrm{v}_{\text{left\_ear}} = \mathrm{k}_{\text{left\_ear}} - \mathrm{k}_{\text{head\_bone}}$$

$$\mathrm{v}_{\text{right\_ear}} = \mathrm{k}_{\text{right\_ear}} - \mathrm{k}_{\text{head\_bone}}$$

where $\mathbf{k}_{\text{head\_bone}}$ denotes the 3-D positions of the 3-D mouse model's skeleton bone head_bone. Under the constraints of anatomy and biomechanics, the weighted addition of these two vectors and the current orientation of head_bone (controlled by NURBS curve) constitutes the new orientation of head_bone, which is mainly the rotation of its local coordinate system around the bone extension direction (Z axis). The rotation of head_bone drives the rotation of the upper body to adapt to the head orientation through inverse kinematics with constraints and weights.

**Supporting Information 2: Semantic Loss**

The fine-grained semantic loss L-semantic is computed as a weighted combination of pixel-level losses across the entire image:

$$L_{semantic} = \sum_{p \in \Omega} w(p) \cdot L_{pixel}(p)$$

where:

- Ω represents the set of all pixels in the image.
- $L_{pixel}(p)$ is the per-pixel loss at pixel p.
- $w(p)$ is the dynamic semantic weight assigned to pixel p, defined in terms of semantic categories (mouse parts or background).

The weight $w(p)$ incorporates semantic information, balancing the importance of:

1. Mouse vs. Background: Assigning equal total weight to pixels belonging to the mouse and background, irrespective of their relative areas.
2. Body Parts vs. Regions: Ensuring smaller body parts (e.g., ears, tail) are given equal consideration as larger body regions (e.g., torso).

$$w(p) = \frac{w_{region}(p)}{A_{class}}$$

where:

- $A_{class}$ is the total area of the semantic class (e.g., background, mouse part) to which pixel $p$ belongs.
- $w_{region}(p)$ assigns a region-specific weight, computed as:

$$w_{region}(p) = \frac{\alpha}{N_{region}}$$

where:

  - $\alpha$ is a balancing factor (e.g., ensuring equal total weight for the mouse and background).

- $N_{region}$ is the number of distinct regions (e.g., body parts, background).

To ensure equal total weight for different body parts, I further assign higher weights to smaller body regions $r$ and lower weights to larger ones such that:

$$w_{region}(p) = \frac{\alpha}{N_{region}} \cdot \frac{\beta_{region}}{A_{region}}$$

where:

- $\beta_{region}$ is the predefined importance for a specific region (default 1).
- $A_{region}$ is the area of such a region.

To ensure equal total weight for mouse and background:

$$\sum_{p \in \Omega_{mouse}} w(p) = \sum_{p \in \Omega_{background}} w(p)$$

where $\Omega_{mouse}$ and $\Omega_{background}$ denote the pixels corresponding to the mouse and background, respectively. This balance is achieved by normalizing the weights within each semantic class.

Combining all the above components, the fine-grained semantic loss becomes:

$$L_{semantic} = \sum_{p \in \Omega} \frac{\alpha \cdot \beta_{region(p)}}{N_{region} \cdot A_{class(p)}} \cdot L_{pixel}(p)$$

In low-resolution, large-scene open field settings, where mice occupy very few pixels and detailed limb features are difficult to discern, a more practical and simplified approach involves balancing the weights inside and outside the mouse's contour at the pixel level. This can be described mathematically as follows:

$$L = \frac{1}{|\Omega|} \left( \sum_{p \in \Omega_{mouse}} w_{mouse} \cdot L_{pixel}(p) + \sum_{p \in \Omega_{background}} w_{background} \cdot L_{pixel}(p) \right)$$

where:

- $\Omega$ denotes the set of all pixels in the image.
- $\Omega_{mouse} \in \Omega$ denotes the set of all pixels in the image.
- $\Omega_{background} = {}^{\Omega_{mouse}}/_{\Omega}$ denotes the set of pixels inside the mouse's contour.

**Supporting Information 3: Feature Engineering**

PyMouse Lifter derives 74 per-frame features from synchronized video and 3-D key-point tracks of freely moving mice. Features fall into three functional families:

| **Family** | **Features** | **Purpose** |
|---|---|---|
| **A. Appearance-change (2-D)** | 9 scalars → **pixel_change** (whole frame) + one 20-px-ROI measure around each of the eight anatomical key points (excluding the two on the tail) | Captures instantaneous intensity change, a hardware-agnostic proxy for body motion or lighting transients. |
| **B. Posture & Kinematics (3-D)** | 24 aligned coordinates + 15 spine-segment kinematic scalars (angle, angular velocity, angular acceleration × 5 segments) + 1 global **bend ratio** | Quantifies absolute body shape after translating the tail root to the origin and rotating the lumbar–tail vector onto the global **+Y** axis. |
| **C. Locomotor dynamics (3-D)** | 2 × (velocity $^{\rightarrow}$, acceleration $^{\rightarrow}$, speed) of **head** and **tail root** (14 scalars) + rolling 8-frame standard deviations of spine angular velocity (5) and head/tail velocity $^{\rightarrow}$ (6) | Measures how rapidly the animal moves through space and the short-term variability of that motion. |

All 3-D positions are expressed in millimeters, then translated so that tail_base = (0,0,0) and rotated about Z such that the tail_base → lumbar_spine vector points along +Y. This alignment removes inter-frame yaw and global translation, ensuring that downstream models learn posture rather than camera orientation.

Quantities requiring history are set to NaN for the first 7 frames; these frames are dropped before flattening so that exported feature tables contain no undefined values.

After flattening, each feature becomes a column named:

- k_3d_x, k_3d_y, k_3d_z for aligned coordinates (k ∈ {nose,…,tail_base});
- segment_angles_0 … segment_ang_acc_4, etc., for spine metrics;
- single-value metrics retain their base name (bend_ratio, pixel_change, …).

The complete Python implementation, together with an extended feature glossary, is hosted at github.com/your-repo/PyMouseLifter (directory features/). The full glossary is also supplied as "PyMouse Lifter Feature Definitions.md" in the repository.

**Supporting Information 4: Depth Estimation Metrics & 3-D Keypoint Errors**

| Training (single-mouse) | Higher is better ↑ | | | Lower is better ↓ | | |
|---|---|---|---|---|---|---|
| | $\alpha_1$ | $\alpha_2$ | $\alpha_3$ | AbsRel | RMSE | RMSE(log) |
| Zero-shot (no re-training) | 0.738 | 0.886 | 0.974 | 0.209 | 51.80 | 0.253 |
| Synthetic data | 1.000 | 1.000 | 1.000 | 0.016 | 4.29 | 0.025 |
| Style-transfer synthetic data | 1.000 | 1.000 | 1.000 | 0.016 | 4.36 | 0.027 |
| Depth-camera data | 1.000 | 1.000 | 1.000 | 0.015 | 4.50 | 0.028 |

Table 1: Single-mouse depth-estimation benchmark (mean values).

| Training (dual-mouse) | Higher is better ↑ | | | Lower is better ↓ | | |
|---|---|---|---|---|---|---|
| | $\alpha_1$ | $\alpha_2$ | $\alpha_3$ | AbsRel | RMSE | RMSE(log) |
| Zero-shot (no re-training) | 0.772 | 0.965 | 0.993 | 0.162 | 40.89 | 0.197 |
| Synthetic data | 1.000 | 1.000 | 1.000 | 0.024 | 6.77 | 0.039 |
| Style-transfer synthetic data | 1.000 | 1.000 | 1.000 | 0.017 | 4.88 | 0.027 |
| Depth-camera data | 0.999 | 1.000 | 1.000 | 0.016 | 5.78 | 0.035 |

Table 2: Dual-mouse depth-estimation benchmark (mean values).

Validation of MDE models trained on real / rendered synthetic data / style-transferred synthetic data on actual depth camera measurements. The predicted depth map is aligned with the actual depth map at the ground level, and only the pixels belonging to the mouse are identified and used for calculation. 1/2/3: Fraction of pixels that are within a scale factor of 1.25 to the 1/2/3 power. AbsRel: Absolute relative (in percentage) error. RMSE: Root mean squared error (in meters). RMSE(log): Root mean squared error on the log scale. (A) Metrics computed on our single-mouse openfield dataset (22 mice, each with 1-3 10-min videos). (B) Metrics computed on MoSeq two-mice openfield dataset (2 mice in a 1.5-hour video). Note: the unit for our computation is mm, while for most computer vision tasks the unit is meters.

**Supporting Information 5: 3-D V.S. 2-D Behavior Classification**

| 3D Lifting XGBoost Classification | Labeled Data | | Sensitivity | F1 Score |
|---|---|---|---|---|
| Behavior Categories | Positive | Negative | ± STD | ± STD |
| Grooming | 2136 | 63315 | 0.860 ± 0.051 | 0.652 ± 0.022 |
| Rearing | 9591 | 55860 | 0.926 ± 0.007 | 0.861 ± 0.007 |
| Locomotion | 20786 | 44665 | 0.879 ± 0.005 | 0.889 ± 0.002 |
| Stationary | 32938 | 32513 | 0.878 ± 0.004 | 0.911 ± 0.002 |
| Overall | N/A | N/A | 0.886 ± 0.012 | 0.828 ± 0.006 |

| 2D Only XGBoost Classification | Labeled Data | | Sensitivity | F1 Score |
|---|---|---|---|---|
| Behavior Categories | Positive | Negative | ± STD | ± STD |
| Grooming | 2136 | 63315 | 0.687 ± 0.039 | 0.513 ± 0.031 |
| Rearing | 9591 | 55860 | 0.778 ± 0.017 | 0.685 ± 0.006 |
| Locomotion | 20786 | 44665 | 0.823 ± 0.010 | 0.857 ± 0.006 |
| Stationary | 32938 | 32513 | 0.836 ± 0.006 | 0.868 ± 0.003 |
| Overall | N/A | N/A | 0.781 ± 0.009 | 0.731 ± 0.010 |

Performance comparison of XGBoost classifiers trained with 3-D lifting versus 2-D input. The upper block reports results for models that use 3-D keypoints obtained via monocular depth lifting; the lower block shows models that rely solely on 2-D top-view keypoints. Sensitivity (recall) and F1 score are presented as mean ± standard deviation across the five test folds. Because a separate yes/no classifier was trained for each behaviour, the positive and negative counts indicate the number of labeled frames that do or do not belong to the corresponding class, respectively.

**Supporting Information 6: Speed of Realtime 3-D Behavior Classification**

| Platform | CPU | GPU (power) | Resolution (resize to) | Batch Size | Vram Usage (G) | FPS |
|---|---|---|---|---|---|---|
| WSL | AMD EPYC 7313 | RTX 6000 Ada (300W) | 480X480 | 2 | 2.2 | 33.743 ± 1.685 |
| WSL | AMD EPYC 7313 | RTX 6000 Ada (300W) | 480X480 | 4 | 2.8 | 42.620 ± 3.107 |
| WSL | AMD EPYC 7313 | RTX 6000 Ada (300W) | 480X480 | 8 | 4.0 | 45.119 ± 3.518 |
| WSL | AMD Ryzen 9 9950X | RTX 4000 SFF Ada (70W) | 480X480 | 2 | 2.2 | 18.318 ± 2.062 |
| WSL | AMD Ryzen 9 9950X | RTX 4000 SFF Ada (70W) | 480X480 | 4 | 2.8 | 20.081 ± 1.144 |
| WSL | AMD Ryzen 9 9950X | RTX 4000 SFF Ada (70W) | 480X480 | 8 | 4.0 | 19.572 ± 1.173 |
| Linux | Intel Core i9 13900KF | RTX 4090 (450W) | 480X480 | 2 | 2.2 | 55.798 ± 3.610 |
| Linux | Intel Core i9 13900KF | RTX 4090 (450W) | 480X480 | 4 | 2.8 | 63.240 ± 6.956 |
| Linux | Intel Core i9 13900KF | RTX 4090 (450W) | 480X480 | 8 | 4.0 | 66.680 ± 3.925 |
| Linux | Intel Core i9 13900 | RTX 3090 Ti (450W) | 480X480 | 2 | 2.2 | 30.712 ± 1.182 |
| Linux | Intel Core | RTX 3090 | 480X480 | 4 | 2.8 | 33.035 |

| | | | | | | |
|---|---|---|---|---|---|---|
| | i9 13900 | Ti (450W) | | | | ± 1.427 |
| Linux | Intel Core i9 13900 | RTX 3090 Ti (450W) | 480X480 | 8 | 4.0 | 36.421 ± 1.997 |

Inference speed using the PyMouseLifter pipeline with a lightened XGBoost model, YOLO11m-pose, and DINO-V2-Large as the encoder for Depth Anything in fp16 precision. Batch Size: The number of images sent to video memory for inference simultaneously. A larger batch size incurs an extra delay of [(batch size - 1) × (time per frame)], but it reduces the data transmission bottleneck. FPS: Frames Per Second, reported as mean ± standard deviation. For this analysis we used pre-recorded data as it is not possible to compare different hardware configurations in a truly live capacity. Rates are expected to be similar for a live configuration.

### Supporting Information 7: Rendered Pose V.S. Groundtruth in Reference

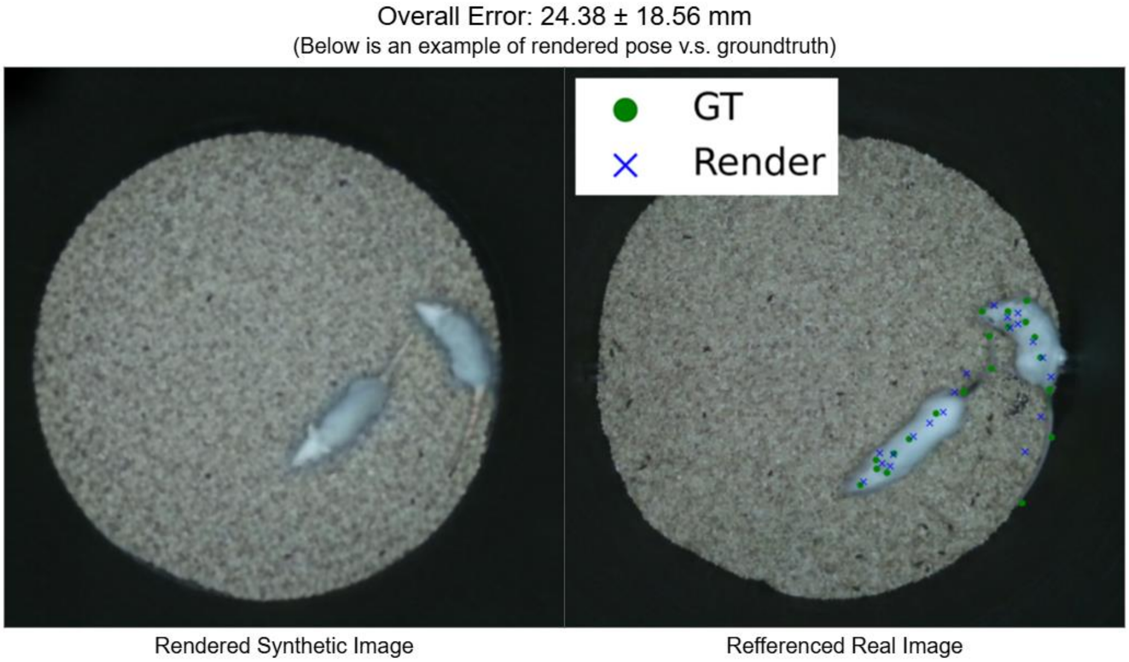


Visualization of the human-labeled groundtruth keypoints in a referenced real image v.s. the pseudo label keypoints from the pipeline after 3-D mouse model optimization. The pose of the 3-D mouse model is reasonable but not exactly the same as the pose of the real mouse, with an average 3-D error of 24.38 ± 18.56 mm.

### Supporting Information 8: YOLO V.S. DeepLabCut in Projection Errors

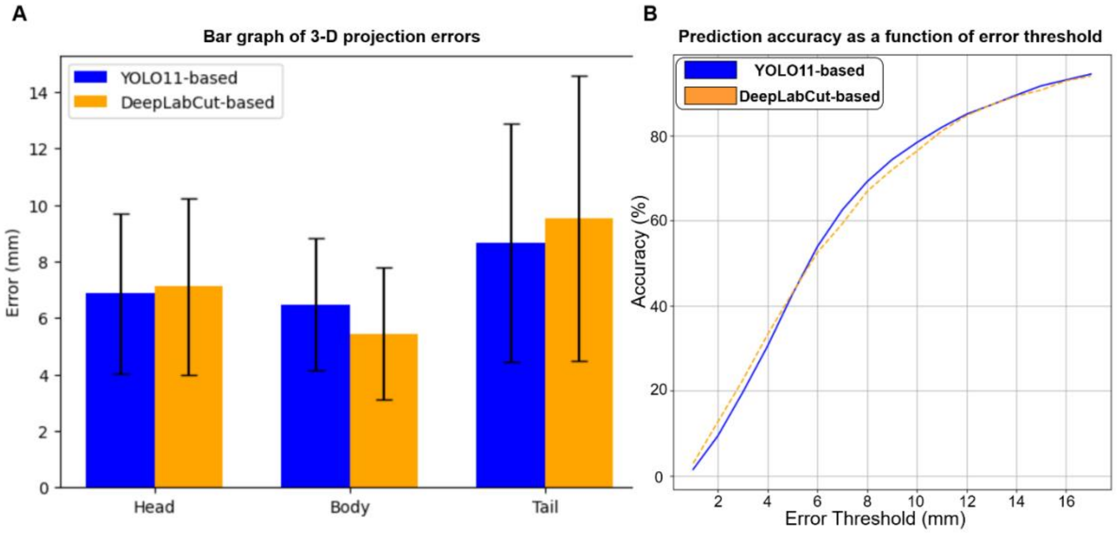


Comparison of the 3-D projection errors using YOLO11m-pose or DeepLabCut SuperAnimal-Topview (Ye et al., 2024) with resnet-50 backbone. (A) Bar graph of 3-D projection errors presented as mean ± SD. (B) 3-D keypoints prediction accuracy as a function of error threshold, for the same data and methods as in (A).
